# Biochemical state in tissue can be detected through ultrasound signal

**DOI:** 10.1101/2024.12.27.630453

**Authors:** Kazuyo Ito, Yuta Iijima, Tomoki Misumi, Gen Hayase, Kazuki Tamura, Kenji Ikushima, Daisuke Yoshino

**Affiliations:** Department of Biomedical Engineering, Graduate School of Engineering, Tokyo University of Agriculture and Technology, Tokyo, Japan; Division of Advanced Applied Physics, Institute of Engineering, Tokyo University of Agriculture and Technology, Tokyo, Japan; Department of Food and Energy Systems Science, Graduate School of Bio-Applications and Systems Engineering, Tokyo University of Agriculture and Technology, Tokyo, Japan; Department of Industrial Technology and Innovation, Graduate School of Engineering, Tokyo University of Agriculture and Technology, Tokyo, Japan; Reseach Center for Electronic and Optical Materials, National Institute for Materials Science, Ibaraki, Japan; Institute of Photonics Medicine, Hamamatsu University School of Medicine, Shizuoka, Japan

## Abstract

Three-dimensional (3D) cell cultures, such as spheroids, are indispensable models for investigating cellular behaviors and responses under conditions that closely resemble in vivo environments. Conventional imaging techniques, including optical microscopy, are often limited by penetration depth and phototoxicity, complicating the analysis of structural and biochemical changes within dense 3D systems. This study demonstrates the application of ultrasound imaging for the non-invasive evaluation of internal dynamics in cancer spheroids over a 15-day period. Scattering-based acoustic parameters revealed spatial variations in brightness and density, correlating with cellular proliferation, apoptosis, and necrosis. Brightness values in central regions progressively decreased after Day 3, approaching near-zero by Day 15, reflecting necrotic core formation. Artificial inhibition of myosin contractility significantly influenced these patterns, providing insights into biomechanical contributions to spheroid organization. The findings establish ultrasound imaging as a label-free, high-penetration technique capable of addressing critical challenges in 3D culture analysis, offering new opportunities for studying cellular dynamics and therapeutic responses in spheroids and organoid models.

## Introduction

The importance of 3D cell culture lies in its ability to closely emulate in vivo environment. Traditional 2D cell culture systems, where cells are grown on flat, rigid surfaces, fail to replicate the complex structural and biochemical characteristics of natural tissue environments. Consequently, cellular behaviors, such as gene expression, metabolism, and drug responses, often diverge significantly from in vivo results, leading to limited clinical relevance ^1–3^. In contrast, 3D cell culture systems, like spheroids and organoids, allow cells to interact in all dimensions, forming cell-to-cell and cell-to-matrix contacts that are essential for maintaining tissue-specific functions and accurately reflecting the microenvironment ^4,5^.

Imaging has a key role in unveiling the complexities cellular dynamics three-dimensionally. However, unlike traditional 2D cultures, which are relatively easy to image, 3D models such as spheroids and organoids are thicker and more structurally complex, and therefore the current imaging technologies have several inherent limitations^6,7^. The challenges associated with optical-based 3D imaging techniques often stem from limitations in depth of field (DOF)^8^. DOF is primarily determined by the wavelength of the light source and the numerical aperture (NA) of the objective lens (i.e. DOF=λ/NA). In confocal laser microscopy, for instance, a mid-range objective lens (NA 0.65, wavelength 633 nm) provides lateral resolution of approximately 600 nm and a DOF of about 2 µm. When using a low-magnification lens (NA 0.2), the lateral resolution decreases to approximately 2 µm, but the DOF extends to around 10 µm. Additionally, specialized pinhole adjustments are employed to achieve an increased DOF with reducing out-of-focus light, leading to slow data acquisition although the pinhole size and DOF are still in a trade-off. Optical Coherence Tomography (OCT) offers greater depth penetration; for example, a swept-source OCT (SS-OCT) system with a central wavelength of 800 nm achieves an axial resolution of 2–5 µm, lateral resolution of approximately 10 µm, and a DOF of around 200 µm^9^. Additionally, phototoxicity and photobleaching are significant concerns in 3D imaging of live cells, as prolonged exposure to high-intensity light can damage cells and degrade fluorescent dyes, impacting the validity of live-cell observations^10^. More recently, spinning disk confocal systems and high-intensity laser-based imaging tools can partially mitigate these challenges by reducing exposure time and improving penetration. They still struggle with complex data processing and large image files typical of high-content imaging in 3D cultures. OCT could be a powerful candidate for visualizing internal dynamics of spheroids^11^. With a combination of OCT and OCT elastography, they can provide mechanical properties of spheroids in 3D^12^. However, limited penetration depth with sub-mm scale and limited holder solutions pose challenges for detailed analysis specifically in real time manner^6^.

Ultrasound (US) could be a suitable alternative because of its low-cost, label-free, and non-invasive imaging capability. It provides two-dimensional cross-sectional images, allowing real-time observation of the shape, boundaries, and internal structures of organs and tissues. It is frequently used for assessing fetal health and diagnosing abdominal organs such as the liver, kidneys, and gallbladder. Expanding applications from clinical to basic biological research, ultrasound technology has a great possibility to observe the internal changes occurring in the tissue or even within the cell. Ultrasonic imaging of spheroids possesses the significant advantage of being able to penetrate deep into the sample, allowing visualization of the internal structure of spheroids on the mm scale. Unlike optical-based imaging there are no limitations in the holding medium as long as the spheroid is anchored at the specific location. Another benefit of using ultrasound is the ability to conduct label-free observations without staining and fixation, thereby preserving the natural state of the cells and tissues under examination. This non-invasive approach minimizes the risk of altering cellular structures and functions, which is a common concern with traditional labeling and fixation methods. Indeed, some researchers succeeded in monitoring cell dynamics through ultrasound. Hagiwara *et al.* visualized changes in ultrasound amplitude with TGF-β1 stimulation to the living fibroblast cells^13^. Another report showed time-dependent changes in ultrasound absorption during mechanical stimulation to the cell nucleus^14^. The advantages of ultrasound, as demonstrated thus far, are particularly maximized in research involving three-dimensional cultured spheroids and organoids. In fact, the initial study conducted by Sherar successfully visualized the internal structures of tumor spheroids^15^. The resulting micrographs given by 100 MHz ultrasound reveal a striking contrast between the necrotic core and the viable rim of the spheroid, demonstrating the method’s capability to provide tomographic images at depths up to 4 mm in biological specimens. Their study demonstrates the potential of ultrasound as a non-invasive technique for detailed imaging of the internal architecture of living tumor spheroids, paving the way for real-time structural analysis in complex 3D cell models.

This time, we investigate the internal changes of cancer spheroids created using a rapid cancer spheroid production method^16^. This method allows for the efficient creation of uniform and reproducible spheroids, which are essential for consistent and reliable research outcomes. Ultrasound imaging was employed to observe cell-dynamics inside spheroids non-invasively and three-dimensionally. 20-MHz center frequency of ultrasound allows to depict the internal structure of the 2 mm radius of spheroid. This research leverages the amplitude of the ultrasound signal, correlating it with the brightness seen in B-mode ultrasound images. Employing established methods for validation, fluorescent images with immunostaining from the spheroid’s center were also captured to monitor biochemical alterations.

The present article conducted two independent experiments assuming that biochemical changes inside the spheroid trigger the changes in the ultrasound signal; first, monitoring the internal state of the spheroids up to fifteen days post-formation, and second, observing changes in the internal state when actomyosin contractility is artificially inhibited. Blebbistatin (BLB) was used as the inhibitor of actomyosin contractility, and a group treated with DMSO alone was measured to control for the solvent’s cytotoxicity. Previous studies have shown that the contraction of spheroids in the BLB-treated group is weaker compared to other groups. Therefore, we hypothesized that the ultrasound brightness values would follow a similar trend, with the control and DMSO groups showing similar patterns since their actomyosin contractility was not inhibited.We found the ultrasound brightness changes with each successive cultivation days, and this trend matches with the fluorescent microscope observations. Notably, the ultrasound brightness showed dynamic changes starting from day 3, which marks the onset of spheroid contraction, and from day 11, when necrosis initiates at the center of the spheroid. The result also reveals this sequential change were disturbed with the inhibition of actomyosin contractility. These results indicate the potential of ultrasound imaging for the evaluation of internal biochemical changes in the spheroid without any destruction.

## Materials and methods

### Cell culture and spheroid formation

Green fluorescent protein (GFP)-labeled MDA-MB-231 cells (human breast adenocarcinoma cell line, AKR-201, Cell Biolabs, San Diego, CA, USA) were cultured in a 75 cm² flask (VTCF75V, VIOLAMO, AS-ONE, Osaka, Japan) using Dulbecco’s Modified Eagle Medium (DMEM; 31,600-034, Gibco, Thermo Fisher Scientific, Waltham, MA, USA) supplemented with 10% v/v heat-inactivated fetal bovine serum (S1810, Bio-West, Nuaillé, France) and 1% v/v penicillin–streptomycin (15,140-122, Gibco). Upon reaching 90% confluence, the cells were harvested using 0.25% trypsin–EDTA (25,200-072, Gibco) and resuspended in DMEM at a concentration of 5 × 10⁷ cells/mL. Spheroid formation was conducted as per protocols previously established in our work [13]. A cell-suspended collagen solution [4.0 mg/mL; native collagen acidic solution (IAC-50, KOKEN, Tokyo, Japan), 10× DMEM, 10 mM NaHCO₃, 10 mM HEPES–NaOH (pH 7.5), and the cell suspension] was prepared on ice to achieve a final concentration of 5 × 10⁶ cells/mL. This solution was then dispensed onto a superhydrophobic multiwell plate and incubated at 37 °C in a 100% humidified atmosphere with 5% CO₂ for 30–60 minutes. Following gelation, primary spheroids with diameters of 2 mm were transferred to a 48-well plate (VTC-P48, VIOLAMO).

Spheroids were incubated under three independent conditions: Group 1: control, Group 2: cultured with (S)0(-)0Blebbistatin diluted with dimethyl sulfoxide (DMSO), and Group 3: cultured with DMSO. Spheroids in group 1 were incubated with no treatment. The spheroids were cultured from day 0, the day they were made, until day 15. Group 2 uses (S)-(-)-Blebbistatin (B592500, Toronto Research Chemicals (Toronto, ON, Canada)) to inhibit actomyosin contractility through the blockade of myosin II-dependent cellular processes^17^. To achieve inhibition of actomyosin contractility, MDA-MB-231 spheroids were incubated from day 0 in an experimental medium containing 5 μM blebbistatin, prepared with dimethyl sulfoxide (DMSO; Fujifilm Wako Pure Chemical Corporation, Osaka, Japan). Group 3 was act as a control for the group 2 considering the toxicity of the DMSO to the cell despite its negligible concentration. Spheroids in group 2 and 3 were cultured from day 0 until day 7.

### Ultrasound measurement

We designed a custom ultrasound laboratory scanning system and built it to acquire radio-frequency (RF) echo-signal data in three-dimensional from cultured spheroids. This system allowed spheroids to be scanned over their entire volume. The scanning apparatus featured a single-element, spherically focused, ultrasound transducer (PT20-6-12.7, Toray Engineering, Japan) with a 6.0-mm aperture length and 12.7-mm focal length. The transducer had a center frequency of 20.0 MHz and a -6dB bandwidth that extended from 4.1 MHz to 31.0 MHz. The axial resolution is calculated as 82.5 µm according to the relationship *c* = *fλ* (*c*: speed of sound of the medium, *f*: frequency, *λ*: wavelength). The theoretically predicted beam diameter was 156.5 µm by using *BD* = 1.02*Fc*⁄*fD* (*F*: focal length, *D*: aperture). The 6-dB depth of field was measured to be 1.60 mm extending from 11.58 to 13.18 mm with the planer reflector measurement. The transducer was excited by a pulser/receiver unit (5073PR, Olympus) and RF echo signals were digitized using a 10-bit at a sampling frequency of 208 MHz.

Fig 1a illustrates the scanning apparatus. Each spheroid was placed in a water tank filled with DMEM. In the water tank, the spheroid was placed on the PDMS substrate as an acoustic absorber. Scan vectors were uniformly spaced by 48 µm in X and Y directions across the entire scan volume to acquire complete full-volume 3D data from each spheroid [38,39]. During the experiments, the focal depth was positioned below the apex of the spheroid. All measurement was conducted between 24.0∼26.0 C°. Temperature of DMEM was measured before and after the measurement and used for the future speed of sound compensation.

**Fig. 1.**
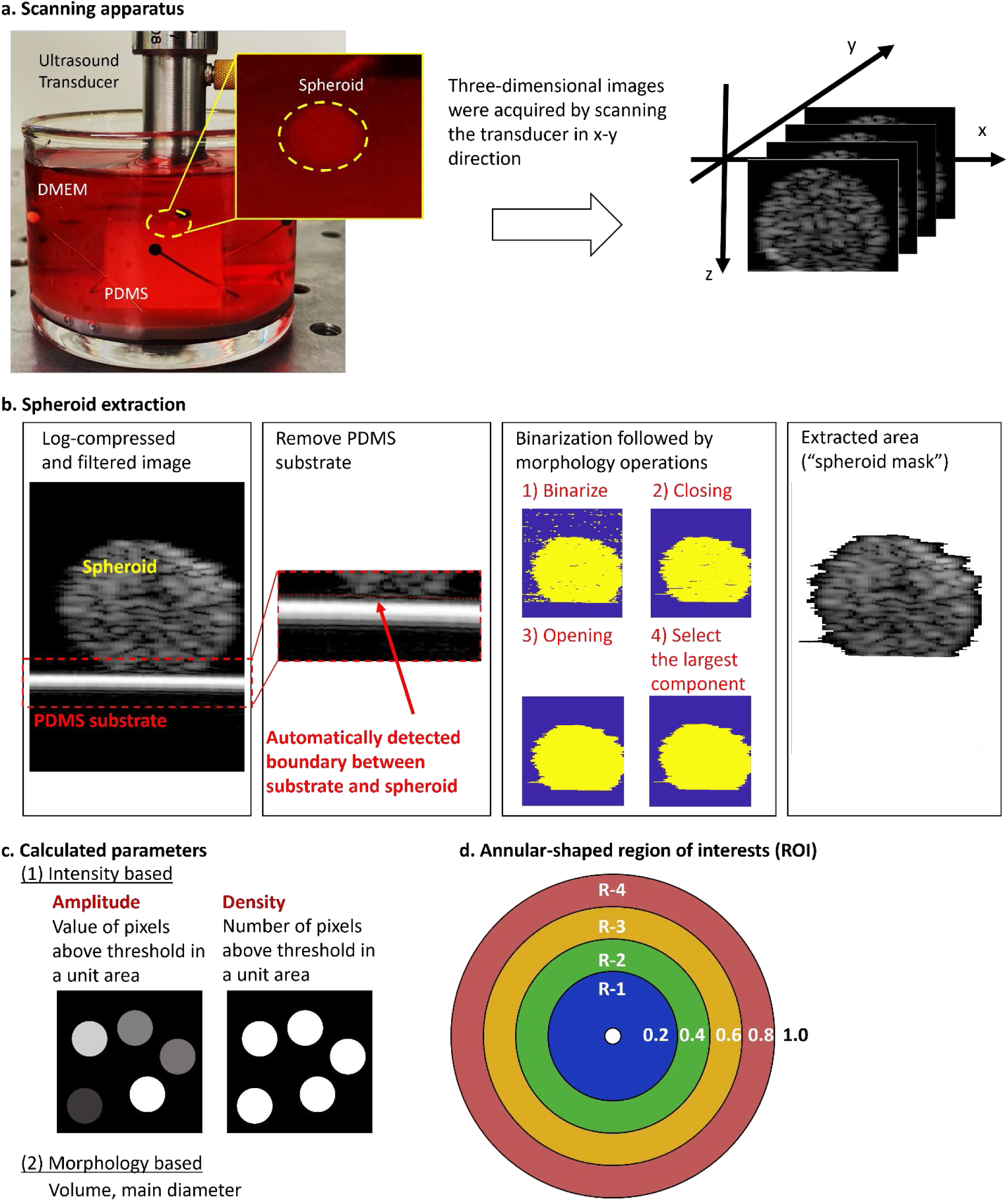
Data acquisition and post-processing. **(a)** Scanning apparatus with a spheroid. By scanning the ultrasound transducer in x-y direction, three-dimensional images were acquired and reconstructed. **(b)** Image-processing to extract spheroid from the image including log compression, PDMS substrate removal, binarization, and morphology operations. **(c)** Overview of the calculated parameters: Intensity-based and morphology-based parameters were calculated. **(d)** Schematic illustration of the annular-shaped region of interest (ROI)s. Taking the long axis of the ellipsoid as 1, the entire spheroid volume is divided into five parts in the radial direction (R-1: inner-most annuli, R-5: outer-most annuli). R-5 was removed because this marginal region possibly contains non-spheroidal information.

### Post-processing

Post-processing can be grouped into three steps: spheroid extraction, parameter calculation, and data summarization based on the distance from the center of the spheroid. Note that the evaluation of the signals was conducted using voltage data, while segmentation analysis was performed on log-compressed data to ensure an appropriate dynamic range. This approach allowed for accurate quantification of signal amplitude while maintaining reliable segmentation of the spheroid regions.

Acquired 3D radio-frequency signal were converted to voltage, followed by the filterization with a 0.1 to 35 MHz band-pass filter. This data has internal information from spheroid and was used for the subsequent evaluation (“voltage data”). Independently, to extract the echo signals originating from the spheroid and create a "spheroid mask," a series of signal and image processing steps were performed (Fig.1b). First, the voltage data were log-compressed to ensure an appropriate dynamic range. Next, cross-correlation with a reference signal was applied to identify and remove echo signals originating from the PDMS layer, allowing for isolation of the spheroid region. The image was then binarized by a threshold that was equal to the average image intensity x4. Noise in the binary image that remained after thresholding was removed by opening and closing morphological operations using disk-shaped structuring element with a radius of 3 pixels. To remove other unwanted segments such as the anterior segment, only the largest connected component in the thresholded image was retained. Morphological operations were conducted to eliminate residual noise and refine the extracted spheroid mask.

For the regions identified as spheroids, two quantitative parameters were calculated as indicators (Fig.1c). The first parameter, amplitude, represents the brightness value per unit area, expressed as the voltage signal. The second parameter, density, was defined as the number of pixels per unit area that exceeded the noise level, regardless of their voltage values, and were therefore classified as signal-detected pixels. As morphological features of spheroids, main diameter and volume were calculated by ellipsoid approximation for the extracted spheroid volume data. For this estimated ellipsoid, five annular-shape regions of interest (ROI) were placed as a function of distance from the center (Fig.1d, R-1: inner-most annuli, R-5: outer-most annuli). The boundary region,R-5, was excluded from subsequent analysis to account for the possibility that it may contain non-spheroidal information due to a segmentation error. Mean and standard deviation were calculated within each ROI. All processing was performed in MATLAB 2022a (The MathWorks, Natick, MA, USA) with the Image Processing Toolbox and Optimization Toolbox.

Independent of the acoustical evaluation, the feature extraction procedure was applied to the fluorescence microscope image but performed with the ImageJ plugin. The algorithm was presented in great detail in our previous paper^18^. To provide a fair comparison between the different treatment, the brightness of ultrasound and fluorescent microscopy images were both normalized by the control Day-1 from the control group.

### Immunohistochemistry

Cultured MDA-MB-231 spheroids were fixed with 4% paraformaldehyde phosphate buffer saline (PFA; 163–20,145, Fujifilm Wako Pure Chemical Corporation, Osaka, Japan) for 3 h at 4 °C. The spheroids were cryoprotected by soaking in 20 w/v% sucrose/phosphate buffered saline (PBS; 05913, Nissui Pharmaceutical, Tokyo, Japan) for 5 h and 30 w/v% sucrose/PBS for an additional overnight at 4 °C. Fixed spheroids were frozen in optimal cutting temperature compound (45,833, Sakura Finetek Japan, Tokyo, Japan) and cut into 10 or 15 μm-thick frozen sections on cryofilm using a cryostat (CM3050S or CM1860; Leica Microsystems, Wetzlar, Germany). After cutting out the frozen sections, the MDAMB-231 cells were permeabilized with 0.1% Triton X-100 (17–1315-01, Pharmacia Biotech, Uppsala, Sweden) in Trisbuffered saline (TBS), followed by incubation in 1% Block Ace (BA; UKB40, DS Pharma Biomedical, Osaka, Japan) in TBS to prevent nonspecific antibody absorption. The cells were then stained using the primary and secondary antibodies diluted in 1% BA in PBS and PBS, respectively, at predefined concentrations (Supplementary Table 1 and 2). Cell nuclei and actin cytoskeleton were stained using 4ʹ,6-diamidino-2-phenylindole dihydrochloride (DAPI; D1306, Invitrogen, Thermo Fisher Scientific) and Alexa Fluor 633 phalloidin (A22284, Invitrogen, Thermo Fisher Scientific), respectively. Stained MDA-MB-231 spheroid sections were observed using a fluorescence imaging system (THUNDER Imaging System, Leica Microsystems, Wetzlar, Germany).

### Experimental reproducibility

All values are shown as mean ± standard deviation (SD) unless stated otherwise. Each data was obtained from three independently repeated experiments.

## Results

### Ultrasound has an ability of MDA-MB-231 cell dynamics observation within the fast fabricated spheroid

Figure 2 demonstrates the ability of ultrasound to observe the dynamics of MDA-MB-231 cells within the rapidly fabricated spheroid. Figure 2a illustrates the capability of ultrasound to inspect the internal changes of the spheroid in three dimensions. Figure 2b shows that during the early phase (days 1 and 2), the spheroid exhibited a homogeneous but sparse scatterer distribution. From days 3 to 9, as cellular density increased, the ultrasound scatter became densely packed. From day 11 onwards, a dense outer shell was observed, indicating the formation of a necrotic core. Figure 2c presents ultrasound-based volume changes with culture duration. Each spheroid exhibited various features depending on the cultured day. Rapid contraction of spheroids began on day 2 after fabrication (2.58±0.30 mm^3^ at day 1 and 2.17±0.20 mm^3^ at day2) and relaxed after day 3 (1.25±0.11 mm^3^), with the volume gradually decreasing until day 15 (0.47±0.02 mm^3^) (Fig.2c). Figures 2d and 2e display quantitative data averaged across each region of interest (ROI) located at every 20% distance from the center of the equivalent unit sphere, supporting these insights. During the early phase (days 1 to 2), the amplitude averaged over the spheroid was lower (0.30±0.03 mV for day1 and 0.34±0.03 mV for day2). From day 3 onwards, as the spheroid began to contract, the global amplitude increased drastically (0.50±0.03 mV) and then gradually decreased over time. Region-dependent changes were also observed, with the amplitude decreasing more sharply at the center of the spheroid (R-1, R-2) compared to the periphery, where the decrease was more gradual (Fig.2f). While little difference was seen between the inner and outer areas on Day 1 (0.30±0.04 mV at R-1 (inner-most), and 0.34±0.03 mV at R-4 (outer most)), the difference increased from Day 2 (0.51±0.03 mV at R-1 (inner-most), and 0.60±0.03 mV at R-4 (outer most))and became particularly pronounced from Day 7 onwards (0.27±0.03 mV at R-1 (inner-most), and 0.51±0.03 mV at R-4 (outer most)). In addition, the brightness values in the central areas R1 and R2 decreased rapidly from Day 3 onwards, and by Day 15 the brightness values were almost approaching 0. Conversely, the density did not change drastically over the spheroid until day 11 regardless of the distance from the center (Figs. 2e and 2g). A substantial decrease in density occurred in the later stages of incubation (days 13 and 15), particularly notable at the center of the spheroid (Fig.2g).

**Fig. 2.**
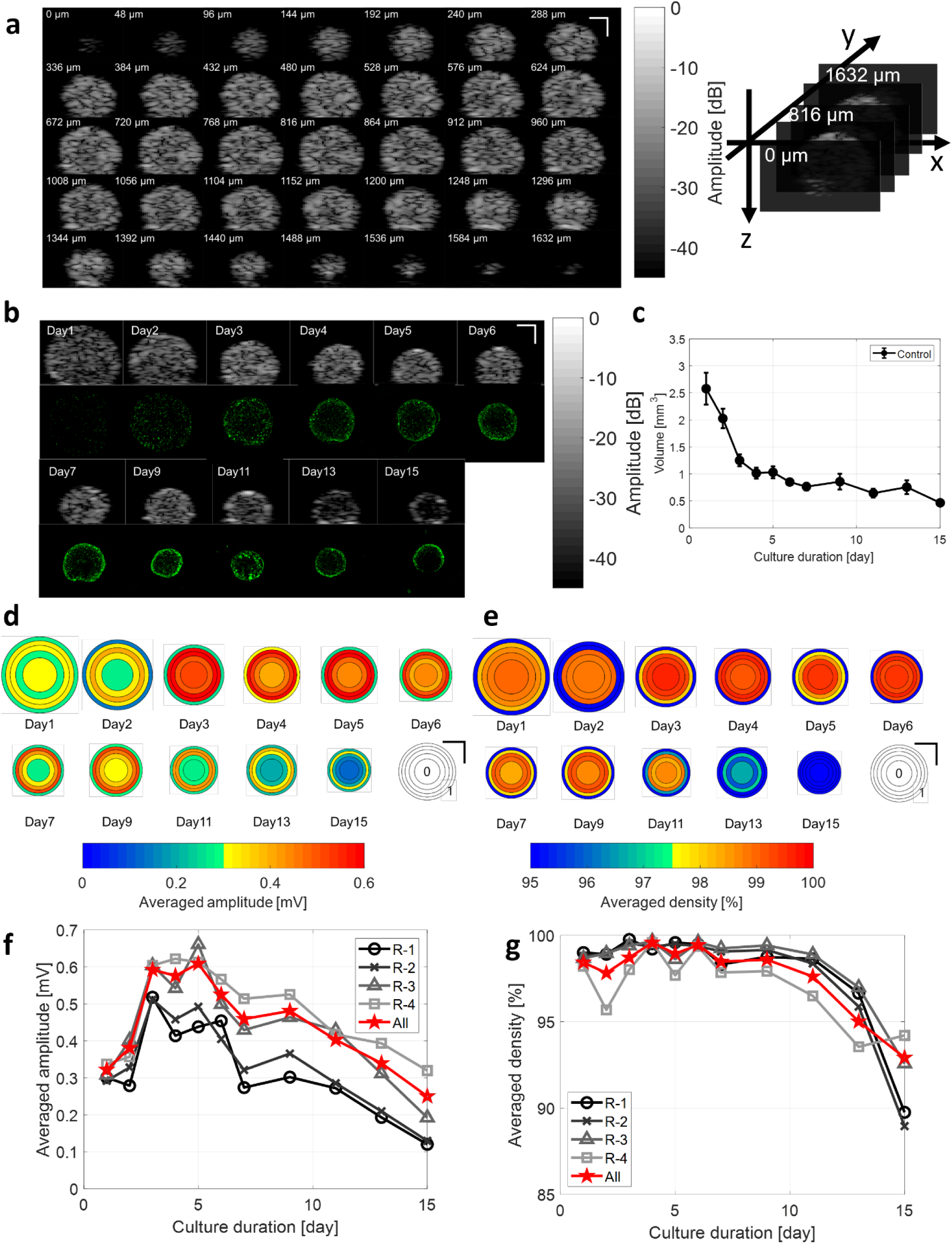
**(a)** Ultrasound brightness (US B-mode) images of the entire spheroid on day 3. The top left image represents the left edge of the spheroid, while the bottom right image represents the right edge. The numbers within the images indicate the position of each respective slice within the entire spheroid. Scale bar=500 μm. **(b)** Time sequence of US B-mode images and GFP fluorescent image at each corresponding day at the center of each spheroid depicting internal changes in the process of its contraction. Each US B-mode image was log-compressed following to the normalization by the amplitude of the PDMS substrate. Scale bar=500 μm. **(c)** Spheroid volume calculated based on the US B-mode images (*n=3*, average±standard deviation in mm^3^). **(d)** Details of changes in the US B-mode images-based amplitude averaged across each region of interest (ROI) located at every 20 % distance from the center of the equivalent unit sphere. The right-bottom panel represents the example of each ROI. Scale bar=500 μm. **(e)** Details of changes in the US B-mode images-based density averaged across each region of interest (ROI) located at every 20 % distance from the center of the equivalent unit sphere. The right-bottom panel represents the example of each ROI. Scale bar=500 μm. Changes in the **(f)** amplitude (in mV) and **(g)** density (in %) of US B-mode images averaged across entire spheroid (All), and averaged across each ROI located at every 20 % distance from the center of the equivalent unit sphere. Circle symbols represent the most-proximal regions (i.e. R-1), cross and triangle symbols represent in between (i.e. R-2 and 3), and rectangle symbols represent the most-distal regions (i.e. R-4). Each dot on the chart represents the mean value, and the whiskers represent the standard deviation.

### Ultrasound unveils the internal changes of spheroid caused by myosin inhibition

Next, we investigated the changes in ultrasound brightness values when actomyosin contractility was artificially inhibited. Figure 3a shows the central cross-section of representative ultrasound B-mode images. As shown in Figure 3, brightness and size change with the number of days of culture. The control and DMSO groups exhibited similar trends. From day 3 onwards, the brightness, particularly in the central region, increased significantly, corresponding to the rapid contraction of the spheroids. In contrast, the BLB-treated group showed similar brightness values to the other groups on days 1 and 2, but these values remained almost unchanged from day 3 onwards. The density showed similar trends across all groups (Figure 3c). Figures 3d to 3g display fluorescent imaging images. DAPI fluorescence was confirmed in the BLB group (Figure 3e). Additionally, as shown in Figures 3f and 3g, the expression of actin and phosphorylated myosin light chain (pMLC) was reduced in the BLB group.

**Fig. 3.**
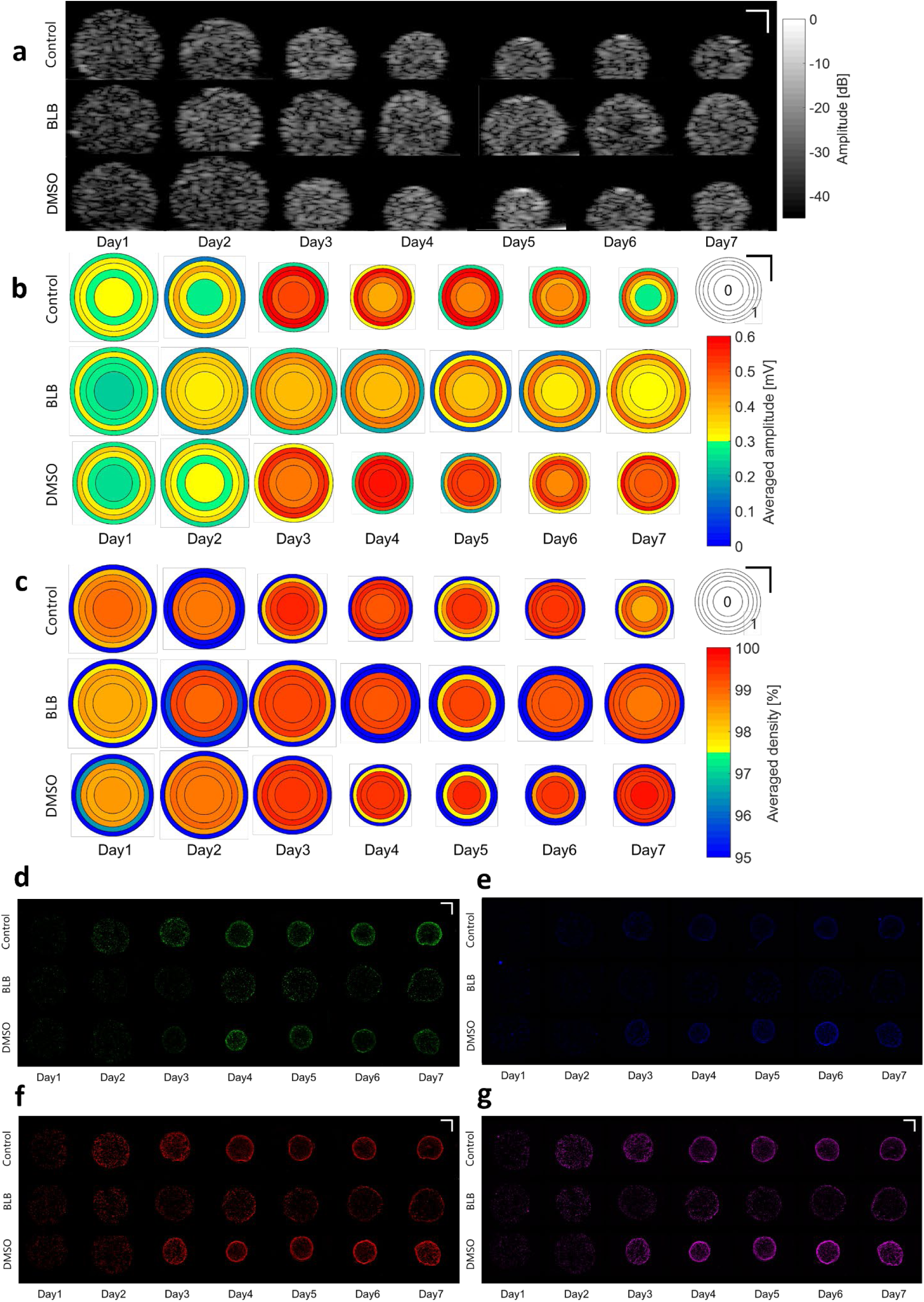
**(a)** Time sequence US B-mode images depicting spheroid contraction under the condition of myosin inhibition with blebbistatin (BLB) (middle), and those without myosin inhibition with dimethyl sulfoxide (DMSO) (bottom). Top row represents control for comparison. Each US B-mode image was log-compressed following to the normalization by the amplitude of the PDMS substrate. Scale bar=500 μm. **(b)** Details of changes in the US B-mode images-based amplitude averaged across each region of interest (ROI) located at every 20 % distance from the center of the equivalent unit sphere. Top row shows the data from control group, middle row depicts the value with BLB, and bottom row depicts the value with DMSO. The top-right panel represents the example of each ROI. Scale bar=500 μm. **(c)** Details of changes in the US B-mode images-based density averaged across each region of interest (ROI) located at every 20 % distance from the center of the equivalent unit sphere. Top row shows the data from control group, middle row depicts the value with BLB, and bottom row depicts the value with DMSO. The top-right panel represents the example of each ROI. Scale bar=500 μm. Localization of GFP **(d)**, DAPI **(e)**, phosphorylation of MLC **(f)**, and F-actin **(g)** in MDA-MB-231 spheroids. Scale bar, 500 µm. From left to right, each column shows each cultured day.

To further elucidate the points demonstrated in Fig. 3, we observed that the expression levels of actin were similar across all three groups until day 2 of culture (Fig.4a). From day 3 onwards, the control and DMSO groups showed an increase in actin expression, followed by a gradual decrease. This increase was particularly steep in the control group. In the BLB group, although actin expression also increased on day 3, the increase was more gradual, and subsequent changes were smaller compared to the other groups (Fig. 4a). The changes in ultrasound brightness values, though exhibiting a smaller dynamic range than actin expression levels, followed a similar trend (Fig. 4b). As observed in the images, density showed an upward trend from day 3 onwards, but there were no significant differences between the groups or across the culture days (Fig. 4c). Interestingly, the DMSO group displayed different results from the control group until day 2. The volume calculated from ultrasound images showed an inverse trend to that of the actin expression and brightness (Figs. 4a and 4b). Although the volume in the BLB group decreased, it did so more gradually compared to the other two groups. The control and DMSO groups exhibited a rapid decrease in volume from day 2 onwards, reaching a plateau after day 4. The scatterplot of volume versus amplitude derived from ultrasound is shown in Fig. 4e. Two distinct groups could be identified: (1) a group with smaller volumes and higher brightness, which included the control and DMSO groups from mid to late culture stages, and (2) a group with larger volumes and medium to low brightness, which included all culture days of the BLB group and the early culture stages of the control and DMSO groups. Finally, we examined the region-dependent changes. While the overall trend was similar across all groups and regions, brightness varied depending on the distance from the center. Notably, in the BLB group, the regions within 50% of the distance from the center showed similar trends regardless of the culture day. The difference in brightness between the outer shell and inner shell was relatively large.

**Fig. 4.**
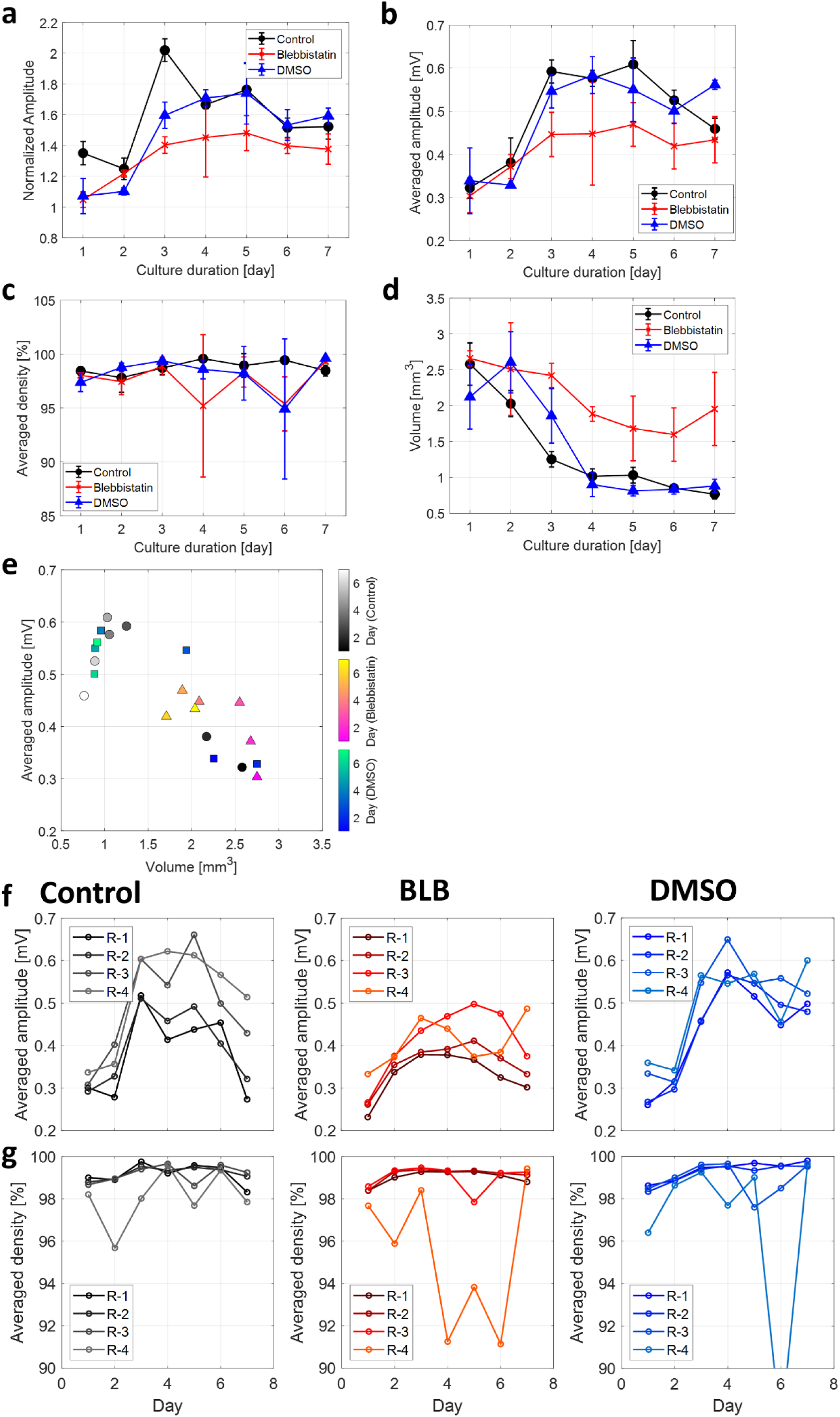
**(a)** Relative expression level phosphorylation of myosin light chain 2 (pMLC) normalized by the intensity of DAPI staining in the 20% region from the center at day1. Changes in the **(b)** amplitude (in mV) and **(c)** density (in %) of US B-mode images averaged across entire spheroid (*n*=3). Both data were normalized by the intensity of US B-mode images in the 20 % region from the center at day 1. **(d)** Spheroid volume calculated based on the US B-mode images. Each dot on the chart represents the mean value, and the error bar represents the standard deviation. Detailed changes in the US B-mode images based **(e)** amplitude (in) and **(f)** density (in %) averaged across each ROI located at every 20 % distance from the center of the equivalent unit sphere. From left to right, each graph represents control, BLB, and DMSO group, respectively. **(g)** Scatterplots comparing US amplitude and US-based spheroid volume (average value, *n=3*). Circle, triangle, and rectangle markers represent control, BLB, and DMSO-treated groups, respectively. The color of the dot indicates the cultured day ranging from day 1 (black for control, magenta for BLB, and blue for DMSO) to day 7 (white for control, yellow for BLB, green for DMSO).

## Discussion

The success of life science experiments often depends on the skill of the experimenter and the use of advanced instruments like confocal laser microscopes and flow cytometers, which require specialized expertise to operate. While these tools enhance our understanding of biological systems, they highlight the need for more accessible technologies that allow non-invasive real-time monitoring. Ultrasound (US) imaging, with its low cost, label-free, and non-invasive capabilities, offers a valuable alternative, particularly for observing three-dimensional cultured spheroids and organoids. This study with 20-MHz ultrasound demonstrated two key findings. Firstly, ultrasound has the capability to observe the internal dynamics of three-dimensional cultured cancer spheroids over time without labels. Secondly, it can differentiate changes in spheroid dynamics when myosin contractility is inhibited by blebbistatin. These findings indicate the effectiveness of ultrasound in monitoring internal changes within cancer spheroids, showing dynamic changes in brightness that correlate with cell contraction and necrosis. Importantly, these findings, which agree with previous reports, were achieved without staining and non-destructively.

### Ultrasound Imaging Unveils Internal Cellular Dynamics in Spheroids

Ultrasound echo images are influenced by various physical phenomena, primarily reflection and scattering, that occur during the propagation of sound waves. Generally, when the echo source is significantly larger than the ultrasound wavelength, reflection occurs, whereas scattering occurs when the echo source is considerably smaller. In the case of the spheroids measured in this study, no scatterers larger than the wavelength exist within the structure, making scattering the dominant source. When scattering is the primary physical phenomenon, the echo signal comprises the summation of backscattered waves from point scatterers within the ultrasound-irradiated region. If point scatterers are densely packed within this region, the backscattered waves from each scatterer interfere with each other, causing the amplitude of the received echo signal to vary significantly depending on the interference state. This interference pattern results in a texture that is uncorrelated with the spatial distribution of the scatterers^19^. The changes in ultrasound backscatter intensity are related to a combination of intrinsic scatterer properties (i.e. acoustic impedance contrast between scatterer source and surrounding medium, scatter location, density, and size of the scatterer) ^20^.

From Figures 2g and I, the results suggest that there are five distinct states throughout the entire culture period. During Days 1-2, the overall brightness and density are low, with minimal spatial variation, indicating State 1. On Day 3, we observe a maximum in amplitude (and possibly density), which corresponds to State 2. Between Days 4-5, although the density remains unchanged, the brightness varies, signifying State 3. During this period, the number of bright pixels remains constant, but the brightness decreases. This suggests reduced interference due to a decrease in the number of scatterers within the resolution cell, as supported by the simulation results of Ohya et. al^19^. From Days 6-11, differences in both density and brightness start to emerge from the inner to outer regions, marking State 4. Finally, on Days 13-15, significant differences in both density and brightness are observed, defining State 5. From Day 3 onwards, despite no significant changes in volume, there are evident changes in brightness and density (Supplementary Fig. 1c, d, e), indicating that significant alterations within the spheroid are occurring, which are not reflected in volumetric changes.

Regarding the biochemical changes during the spheroid culture process, our previous research with protein assays has provided some insights ^18^. As the spheroids mature, the internal cell density increases, leading to hypoxia and subsequent HIF-1α expression. This process is followed by the development of a necrotic core at the center, along with the activation of Caspase-3. They further noted that tumor cells with high proliferative potential (Ki-67 positive cells) migrated to the outer layer of the spheroid as the culture period progressed. Considering these results in conjunction with the actual changes occurring within the spheroids, we can interpret the distinct states as follows: State 1 corresponds to the phase where cells are proliferating. During this phase, the distance between cells is relatively large, meaning the number of scatterers within the resolution cell is low, resulting in low intensity and density. State 2 marks the peak of cell proliferation (when Ki-67 expression is at its maximum), leading to a rapid increase in the number of scatterers within the resolution cell. State 3 occurs when the nutrients and oxygen necessary for cell proliferation can no longer reach the center of the spheroid, causing cells to begin dying from the center. Although these cells are dead, they are not removed, as confirmed by DAPI staining^18^. Consequently, the brightness in the central region decreases, but the density remains largely unchanged. The analysis did not reveal whether there was variability in the size of the cells near the center. State 4 indicates the onset of hypoxia in the center, prompting cells to begin collective migration from the inside to the outside. This results in increased brightness at the periphery and decreased density in the center due to the absence of cells. State 5 is characterized by the formation of a necrotic core (necrosis), where cells swell and rupture. This reduces the acoustic impedance difference with the surrounding area, leading to a decrease in both brightness and density.

The brightness of the ultrasound and the expression levels of actin showed similar trends (Fig. 4a, b), suggesting that the brightness of the ultrasound is related to myosin contractility. This correlation likely arises because the contraction of the spheroid increases the number of cells per unit area, thereby enhancing the number density of ultrasound speckles. This shift alters the interference state of the sound waves, leading to an increase in brightness. Notably, while the amplitude changed, the density remained constant, indicating that cell contractility was modulated without inducing necrosis, consistent with previous studies. The stable density further suggests that the proliferation function is maintained, indicating ongoing cellular activity. After Day 3, the density was slightly lower in the BLB group compared to the control and DMSO groups, likely due to a reduction in the number of scatterers per unit area caused by the inhibition of contractility. Changes in brightness from the center to the periphery followed similar trends as seen in the control group, although this effect was specific to the BLB group and differed from the control and DMSO groups, suggesting that ultrasound brightness could serve as a marker to distinguish between these groups. Moreover, at the two-dimensional single-cell scale, previous studies based on optical tweezers have demonstrated that the addition of blebbistatin leads to changes in mechanical stiffness^21^. Force spectrum microscopy, also based on optical tweezers, has further shown that blebbistatin suppresses the random intracellular forces in the cytoplasm^22^, a finding corroborated by similar trends observed in atomic force microscopy (AFM) analysis^23^. Additionally, OCT-based observations of spheroids have reported that blebbistatin induces cellular relaxation, increasing the fluidity of organelles within the cytoplasm^24^. Collectively, these findings align with our ultrasound data, supporting the notion that changes in actomyosin contractility due to blebbistatin administration can be effectively monitored through ultrasound, offering new insights into cellular dynamics within 3D spheroid models.

### Advantages of Ultrasound Imaging Principles

In recent years, various imaging techniques have been utilized for non-invasive observation of 3D cell culture models, such as spheroids, to assess their structural and functional dynamics^5^. Each method, including Optical Coherence Tomography (OCT), Förster Resonance Energy Transfer (FRET), and fluorescence microscopy, offers unique advantages and limitations that complement ultrasound’s capabilities.

OCT has an ability to illustrate time-dependent dynamics occurring in spheroids^11,24^. At cellular or even sub-cellular imaging was achieved. OCT, a powerful light-based imaging technique, enables high-resolution visualization of internal structures, making it ideal for detailed morphological assessments of 3D cell cultures. Due to its ability to penetrate deeper than traditional optical methods, OCT provides a valuable tool for non-invasive imaging of spheroid structures^11^. FRET is another highly specialized technique that provides insights at the molecular level, detecting energy transfer between closely situated fluorophores to reveal interactions between specific proteins or molecular structures^25^. This technique is especially valuable for studying biochemical processes, such as protein-protein interactions or conformational changes within cells, at a high level of specificity. FRET’s ability to monitor molecular interactions in real time makes it a powerful tool for investigating cellular mechanisms that drive behavior in 3D models. Traditional fluorescence microscopy, like FRET, provides detailed information on cellular structures and protein localization, excelling in surface-level imaging with high specificity and resolution. This technique is widely used for its versatility and adaptability across various cellular and molecular applications. Many optical-based approaches have been proposed recently to capture more advanced information at sub-cellular scale^26^. For example, expanding on OCT, Optical Coherence Elastography (OCE) provides an added dimension by incorporating elasticity measurements, enabling researchers to assess the mechanical properties of 3D cultures, such as stiffness and elasticity variations^12^. These properties are critical for understanding cellular behavior and tissue-like responses, especially in models that simulate tumor microenvironments. OCE allows for localized assessment of stiffness gradients within the sample, providing insights into how mechanical cues influence cellular responses.

Despite their distinct advantages, each technique faces significant limitations when applied to imaging 3D cultured spheroids. A primary drawback shared among these techniques is their limited depth penetration due to light scattering and absorption, which restricts their ability to capture detailed information from deeper layers within dense spheroids. This limitation is especially pronounced in fluorescence microscopy and FRET, where high-resolution imaging is typically confined to surface regions, making deeper structures increasingly difficult to resolve. Additionally, fluorescence-based methods suffer from photobleaching and phototoxicity, complicating long-term, real-time observation of live 3D cultures, as prolonged exposure to excitation light can alter cellular behavior or compromise cell viability^10^. While OCT and OCE offer improved depth penetration and enable non-invasive imaging, their penetration depth is generally limited to a few hundred micrometers, which is insufficient for imaging larger spheroids (up to ∼5 mm in diameter) like the ones developed in our study^16^, or for future organoid models designed to mimic full organ functionality^27^. Furthermore, OCT and OCE typically require spheroids to be embedded in hydrogels or other stabilizing materials for imaging, which complicate dynamic observations of drug-induced changes, as embedded samples may respond differently to treatment than free-floating or actively growing models. As such, fluorescence microscopy and FRET, both confined primarily to two-dimensional imaging due to limited penetration depth, are similarly constrained when applied to thick, layered spheroid cultures. Collectively, these drawbacks hinder the use of these optical-based modalities for comprehensive and non-disruptive monitoring of dynamic processes within large 3D spheroid and organoid cultures. This highlights the need for alternative or complementary imaging approaches that offer greater depth penetration, minimal invasiveness, and high-resolution capabilities, especially as 3D models continue to expand in size and complexity.

Ultrasound imaging offers several advantages that address the limitations encountered with optical-based methods. Ultrasound’s primary strength lies in its ability to achieve deeper penetration, allowing it to image the internal structures of spheroids and other 3D cultures non-invasively and in real time.

Another significant advantage of ultrasound is its label-free nature, which contrasts with fluorescence-based techniques like FRET and conventional fluorescence microscopy that require staining or labeling. For example, Watanabe successfully clarified The mechanism of regulation of myosin II dynamics in vivo by phosphorylation of myosin regulatory light chain (MRLC) with the fluorescence-probe based approach^28^. They successfully demonstrated myosin II dynamics by live imaging of GFP-tagged MRLC, however, dyes and fluorophores can disrupt natural cellular function and lead to phototoxicity or photobleaching with prolonged observation. Ultrasound imaging eliminates these concerns, enabling real-time, long-term monitoring of cellular and structural dynamics without altering cell viability or behavior. This attribute is particularly valuable for tracking gradual changes in cellular organization, proliferation, migration, and apoptosis over extended culture periods or in response to external stimuli, such as drug treatments. Furthermore, while OCT and OCE typically require samples to be embedded in stabilizing matrices like hydrogels, ultrasound allows for imaging of free-floating samples, making it ideal for assessing dynamic responses to drug administration or environmental changes. This flexibility supports studies of pharmacodynamics and cellular responses in a more native, unconfined environment, preserving the natural conditions of spheroids and organoids during observation.

Previous studies using ultrasound have shown that various factors, such as cell size and cell viability, significantly contribute to changes in ultrasound intensity. For instance, at the microscopic scale, Fadhel et al. reported that acoustic impedance, which relates directly to the intensity of the received ultrasound signal, varies with cell size^29^. They also found that acoustic impedance differs between individual cells and cellular clusters, illustrating the impact of cellular structure on ultrasound properties. Additionally, studies have reported that acoustic intensity changes with apoptosis^30^; this trend has been observed even in cells where apoptosis was artificially induced by drugs^31^. For aggregated cells (though not fully developed spheroids), one study demonstrated that the variance in cell size significantly contributed to the intensity of ultrasound echo signals^32^. In our measurements, a decrease in brightness was observed in regions corresponding to necrosis (Fig. 2d, g), aligning with trends identified in earlier studies by Sherar^15^. Taken together, these findings suggest that internal biochemical and structural changes occurring within the cancer spheroid can be effectively captured through ultrasound signals, making it a promising technique for assessing the progression and treatment response in 3D cell models.

### Limitations

One of the primary limitations of ultrasound imaging is its relatively low spatial resolution compared to these optical techniques. While high-frequency ultrasound improves resolution, this enhancement comes at the cost of penetration depth, limiting its applicability for imaging finer subcellular structures. In contrast, OCT and fluorescence microscopy achieve submicron resolution, allowing for detailed visualization of intricate structures within cells and enabling the study of localized molecular changes, which are essential for understanding cellular processes at a high level of detail. The trade-off between resolution and depth is also a limitation in ultrasound imaging, as achieving high resolution requires higher frequencies that reduce penetration depth, especially when imaging larger or more complex 3D samples.

Moreover, while ultrasound’s label-free nature preserves cells in their natural state, it lacks the molecular tracking ability that labeled optical techniques achieve. Fluorescence microscopy, in particular, excels at highlighting specific molecules and structures, which allows for detailed monitoring of dynamic changes, molecular localization, and interactions within the cellular environment. This is particularly advantageous in studies involving specific cellular or molecular targets where direct labeling enhances observation accuracy. Although ultrasound is advantageous for long-term studies due to its non-invasive nature and absence of phototoxicity, it does not offer the same dynamic range of molecular insights as fluorescence-based techniques.

Further analysis including ultrasound scattering-based quantification, which is called quantitative ultrasound (QUS), is potentially able to address some of these limitations by providing additional information on cellular properties such as density, size, and organization^33,34^. Through QUS metrics, ultrasound can offer insights into mechanical and structural characteristics within 3D cell cultures, which may indirectly indicate cellular viability, proliferation, or structural changes in response to treatments. Leveraging ultrasonic scattering as an indicator may allow for the detection of phenomena smaller than the imaging resolution, potentially addressing spatial resolution limitations that have posed challenges in ultrasound diagnostics. By combining QUS with high-resolution, molecularly specific methods, researchers can achieve a more comprehensive view of cellular processes across spatial scales, providing depth and contextual insights that support a holistic understanding of 3D cell cultures and tissue models.

Nevertheless, as demonstrated so far, we were able to reveal the dynamic internal changes associated with spheroid maturation using ultrasound observation in a label-free and non-invasive manner. Previously, these dynamics could only be elucidated through time-consuming and technically demanding analytical methods such as immunostaining and western blotting. Our approach now allows for straightforward observation of these processes.

## Conclusion

In conclusion, our study demonstrated that ultrasound imaging can effectively reveal the dynamic internal changes associated with spheroid maturation in a label-free and non-invasive manner. The brightness of the ultrasound signal correlated with actin expression levels, indicating a relationship with myosin contractility. This relationship is likely due to changes in the interference state of sound waves caused by variations in cell density and contractility within the spheroid. The constant density and variable amplitude suggest that while cell contractility changes, necrosis does not occur, maintaining cellular activity. Notably, the distinct trends in brightness and density changes between the BLB group and the control/DMSO groups highlight the potential of ultrasound imaging to differentiate these conditions. Thus, our findings suggest that ultrasound can be a powerful tool for observing changes in actomyosin contractility induced by blebbistatin, providing a simpler and more accessible method compared to traditional techniques like immunostaining and western blotting.

## Acknowledgement

This study was partly supported by grants from the Nakatani Foundation for Advancement of Measuring Technologies in Biomedical Engineering and the JSPS KAKENHI (No.21K19893) to D.Y.

## Supplementary Figures

**Supplementary Fig. 1.**
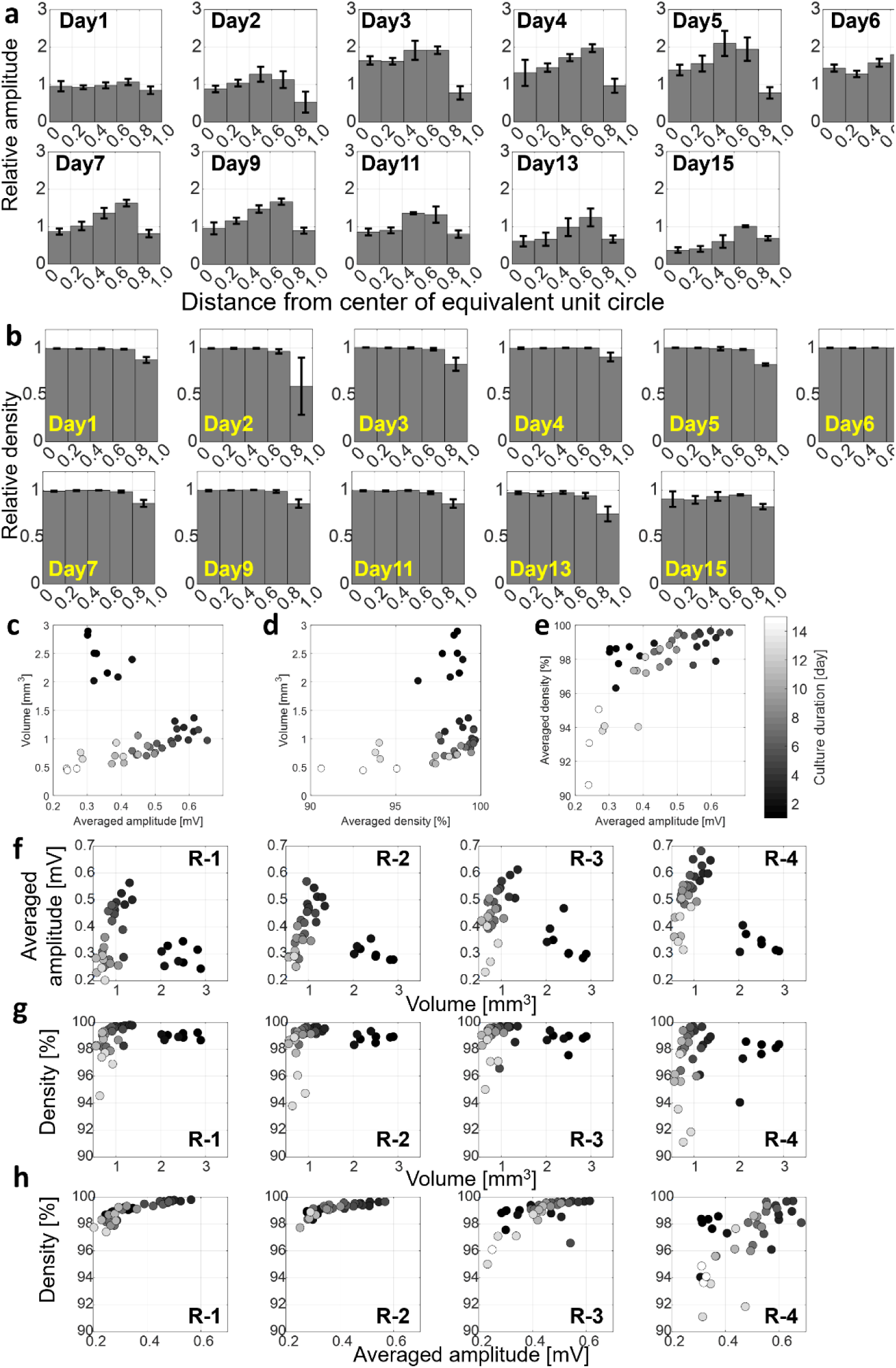
Ultrasound depicts the changes of internal spheroid quantitatively. Average and standard deviation of relative amplitude **(a)** and relative density **(b)** averaged across each ROI located at every 20 % distance from the center of the equivalent unit sphere. After taking average, each data was normalized by the data from the center of day1 (i.e. the center of day1 takes the value of 1). The length of each bar represents the average value and whiskers represents the standard deviation. Scatterplots comparing normalized amplitude (in mV) and ultrasound-based volume (in mm^3^) **(c, f)**, density (in %) and ultrasound-based volume (in mm^3^) **(d, g)**, and density (in %) and normalized amplitude (in mV) **(e, h).** Panels (c) to (e) shows the average across the spheroid, and (f) to (h) shows the breakdowns. From left to right, each panel depicts the most-proximal regions (i.e. R-1), to the most-distal regions (i.e. R-4). Each dot in panel (c) to (h) represents each spheroid. The color of the dot indicates the cultured day ranging from day 1 (darkest, black) to day 15 (lightest, white).

**Supplementary Fig. 2.**
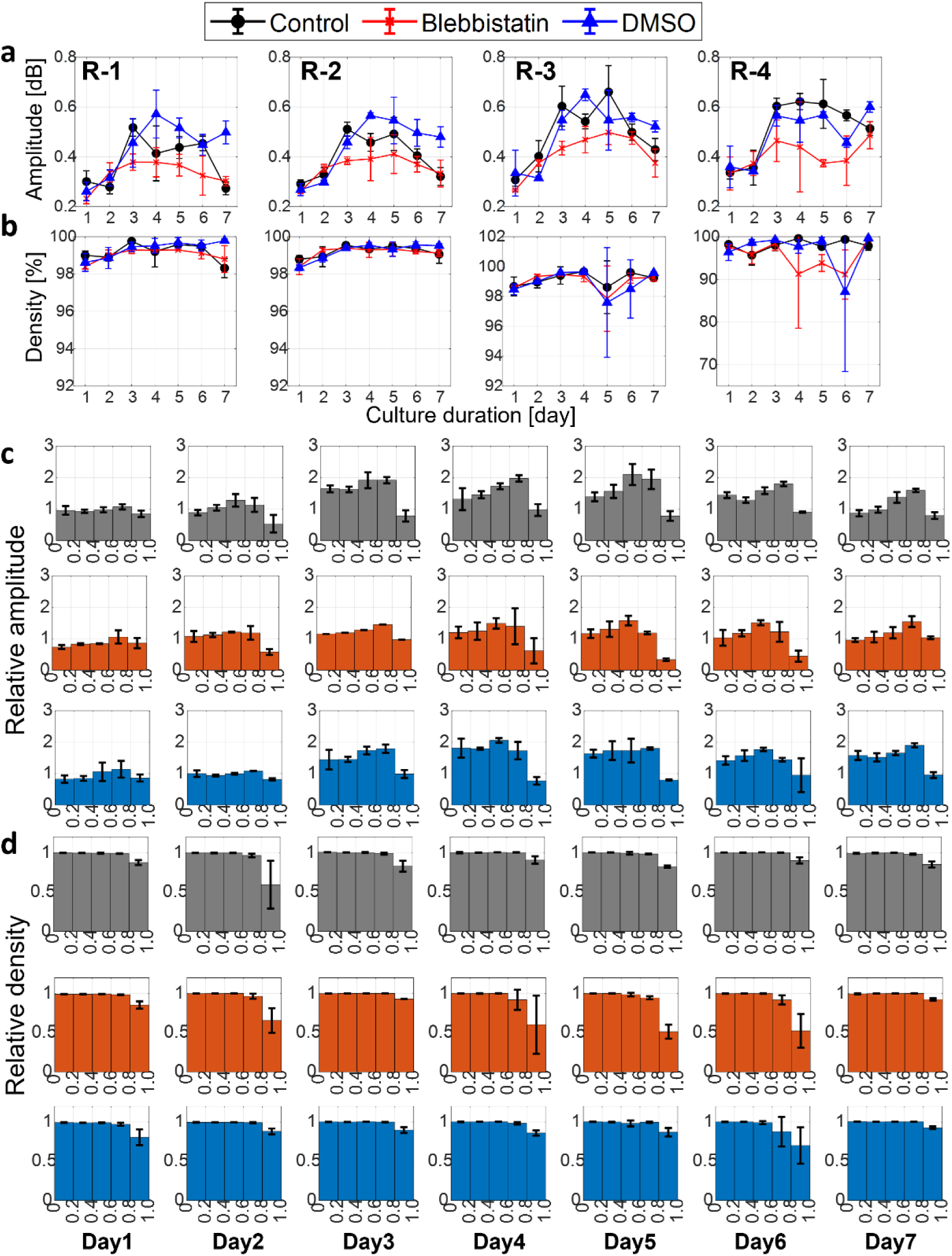
Myosin contractility inhibition causes the changes in the ultrasound brightness. Detailed changes in the US B-mode images based **(a)** amplitude (in mV) and **(b)** density (in %) averaged across each ROI located at every 20 % distance from the center of the equivalent unit sphere. Circle, triangle, and rectangle markers represent control, BLB, and DMSO-treated groups, respectively. Average and standard deviation of relative amplitude **(c)** and relative density **(d)** averaged across each ROI located at every 20 % distance from the center of the equivalent unit sphere. From top to bottom, each row represents control (black), BLB (orange), and DMSO-treated group (blue). After taking average, each data was normalized by the data from the center of day1 (i.e. the center of day1 takes the value of 1). The length of each bar represents the average value and whiskers represents the standard deviation.

## Notes

### Competing Interest Statement

The authors have declared no competing interest.

